# Prostatic osteopontin expression is associated with symptomatic benign prostatic hyperplasia

**DOI:** 10.1101/2019.12.23.887612

**Authors:** Petra Popovics, Wisam N. Awadallah, Sarah Kohrt, Thomas C. Case, Nicole L. Miller, Emily Ricke, Wei Huang, Marisol Ramirez-Solano, Qi Liu, Chad M. Vezina, Robert J. Matusik, William A. Ricke, Magdalena M. Grabowska

## Abstract

**Background:** Male lower urinary tract symptoms (LUTS) occur in more than half of men above 50 years of age. LUTS were traditionally attributed to benign prostatic hyperplasia (BPH) and therefore the clinical terminology often use LUTS and BPH interchangeably. More recently, LUTS were also linked to fibrogenic and inflammatory processes. We tested whether osteopontin (OPN), a pro-inflammatory and pro-fibrotic molecule, is increased in symptomatic BPH. We also tested whether prostate epithelial and stromal cells secrete OPN in response to pro-inflammatory stimuli and identified downstream targets of OPN in prostate stromal cells.

**Methods:** Immunohistochemistry was performed on prostate sections obtained from the transition zone (TZ) of patients who underwent surgery (Holmium laser enucleation of the prostate) to relieve LUTS i.e. surgical BPH (S-BPH) or patients who underwent radical prostatectomy to remove low-grade prostate cancer (incidental BPH, I-BPH). Images of stained tissue sections were captured with a Nuance Multispectral Imaging system and histoscore, as a measure of OPN staining intensity, was determined with inForm software. OPN protein abundance was determined by Western blot. The ability of prostate cells to secrete osteopontin in response to IL-1β and TGF-β1 was determined in stromal (BHPrS-1) and epithelial (NHPrE-1 and BHPrE-1) cells by ELISA. qPCR was used to measure gene expression changes in these cells in response to OPN.

**Results:** OPN immunostaining (p=0.0107) and protein levels were more abundant in S-BPH than I-BPH. Staining was distributed across all cell types with highest levels in epithelial cells. Multiple OPN protein variants were identified in immortalized prostate stromal and epithelial cells. TGF-β1 stimulated OPN secretion by NHPrE-1 cells and both IL-1β and TGF-β1 stimulated OPN secretion by BHPrS-1 cells. Interestingly, recombinant OPN increased the mRNA expression of *CXCL1*, *CXCL2*, *CXCL8*, *PTGS2* and *IL6* in BHPrS-1, but not in epithelial cell lines.

**Conclusions:** OPN is more abundant in prostates of men with S-BPH compared to men with I-BPH. OPN secretion is stimulated by pro-inflammatory cytokines, and OPN acts directly on stromal cells to drive the synthesis of pro-inflammatory mRNAs. Pharmacological manipulation of prostatic OPN may have the potential to reduce LUTS by inhibiting both inflammatory and fibrotic pathways.

## Introduction

Male lower urinary tract symptoms (LUTS) become more frequent with age and are experienced by approximately 60% of men aged 50 and above^1^. LUTS include obstructive (e.g. urinary retention, weak flow and incontinence) and irritative (e.g. urgency, nocturia, dysuria and urinary frequency) symptoms that are associated with a decline in the quality of life^2^. Benign hyperplasia of proliferating epithelial and stromal cells occurs in the transition zone and it is traditionally identified as the leading cause of LUTS. Thus, benign prostatic hyperplasia (BPH) is often used as a synonym of LUTS. The etiology of BPH/LUTS remains poorly understood but multiple factors have been indicated in its pathogenesis, including chronic inflammation that may stimulate proliferation^3, 4^ and trigger periurethral fibrosis^5, 6^. Fibrosis increases tissue stiffness and deposition of extracellular matrix (ECM) components as the result of an unregulated wound-healing process contributing to decreased urethral compliance^6, 7^.

Around 55% of men who visit a urologist because of LUTS are given long-term medical treatments^8^. For symptomatic relief, α1-adrenergic antagonists (a-B) are used to relax smooth muscle cells in the bladder and the urethra^9^ but have no effect on prostatic volume and patients progress to invasive therapy with the same rate as placebo controls ^10^. Although 5α-reductase inhibitors (5ARI) decrease prostate volume^10, 11^, patients may still require surgical intervention due to therapy resistance or side effects including impotence, abnormal ejaculation and fatigue ^10^. Treatment with 5ARIs only reduces glandular tissue whereas the relative abundance of stromal and epithelial cells in BPH nodules is diverse suggesting that this treatment may become less efficient in cases with stromal dominance^12^. Furthermore, both 5ARI and a-B stimulate the TGF-β profibrotic pathway and thus may promote fibrosis^13, 14^. In agreement with this, the MTOPS (Medical Therapy of Prostatic Symptoms) study revealed that fibrosis in the transition zone of the prostate is associated with increased risk of clinical progression of LUTS in patients who received combined a-B and 5ARI treatement^15^. Consequently, more efficient therapies are needed that target not one but multiple causes of LUTS including proliferation, inflammation and fibrosis in the prostate.

We previously showed that carrageenan induces chronic prostatic inflammation in rats and increases mRNA expression of *Spp1,* which encodes osteopontin (OPN)^16^. OPNis a secreted phosphoglycoprotein and an extracellular matrix constituent^17^. There are five OPN splice variants that encode proteins differing in phosphorylation, transglutamination, secretion and potentially, activity^18^. OPN is expressed by most immune cells, is a cytokine and a chemoattractant, and activates T-cells and macrophages^19–21^. OPN has been implicated in the pathogenesis of multiple fibrotic diseases^22–24^. With the new recognition that prostatic collagen accumulation is associated with LUTS severity and treatment resistance^15^, it is critical to identify mechanisms of collagen accumulation, specifically whether OPN participates in prostatic inflammatory and fibrotic pathways. The goals of this study are to: (1) test hypotheses that OPN is more abundant in the prostate transition zone of men with symptomatic BPH compared to that of asymptomatic BPH, (2) determine which prostatic cells synthesize OPN, (3) test whether pro-inflammatory and pro-fibrotic factors increase *OPN* secretion in human prostatic cells, and (4) identify genes that are regulated by OPN inhuman prostatic cells. Our study revealed that OPN is more abundant in men with the symptomatic S-BPH versus incidental BPH levels in multiple cell types. We identified macrophages and prostatic stromal and epithelial cells as sources of OPN and showed that IL-1β and TGF-β1 drive OPN secretion. We also found that OPN increases mRNA expression of proinflammatory *IL-6*, *CXCL1-2* and *-8,* as well as, *PTGS2* in immortalized human prostate stromal cells. These results indicate that OPN may contribute to LUTS by responding to inflammatory signals and amplifying them in the prostate by triggering the expression of cytokines/chemokines in stromal cells.

## Materials and Methods

### Human tissue procurement

Tissue collection was approved by the Institutional Review Board at Vanderbilt University Medical Center (VUMC), and analysis of specimens at Case Western Reserve University (CWRU) and the University of Wisconsin-Madison (UW-M). De-identified prostate tissues were acquired from the VUMC BPH Tissue and Data Biorepository and their acquisition has been described previously^25^. Incidental specimens (n=6) were isolated from patients undergoing radical prostatectomy for low volume, low-grade prostate cancer confined to the peripheral zone of the prostate. Samples were isolated from the transitional zone and were selected from patients who had not taken α-blockers and the malignancy was low risk (Gleason Score 7 or less) and small volume (≤1 cc). Surgical BPH (S-BPH) specimens (n=31) were isolated from patients who failed medical therapy and underwent holmium laser enucleation of the prostate (HoLEP) to relieve LUTS. Inflammation was scored by a pathologist (W.H) and did not significantly differ between I-BPH and S-BPH specimens.

### Immunohistochemistry

Slides were deparaffinized in two changes of xylenes for 5 min. and hydrated in a series of 100%, 90%, 70% and 50% ethanol (v/v%) followed by a washing step in tap water. Antigen retrieval was performed in a citric acid-based solution (Vector H3300) using a microwave (5 min 40 sec with full power and 20 min with 30% of power). Slides were cooled for 20 min, washed in PBS and endogenous peroxidases were blocked with 3% H2O2 in methanol for 20 min. Blocking solution consisted of 1.67% goat serum in PBS and was applied to slides for 30 min at room temperature. Osteopontin (ab8448, Abcam) antibody was diluted 1:500 in blocking solution and applied to slides overnight at 4°C. The next day, slides were washed in PBS and the biotinylated rabbit antibody (1:200 dilution, VECTASTAIN® Elite® ABC Kit, Vector Laboratories) was applied in blocking solution for 1 h. Slides were then washed and the reagent containing avidin and biotinylated HRP was applied to slides for another 60 min. This was followed by 3 washing steps in PBS and the signal was developed with ImmPACT® DAB Peroxidase Substrate (SK-4105, Vector) timed under a microscope. Mouse spleen and kidney were used as positive controls to set the exposure time for human specimens. Development was terminated in tap water and nuclei were counterstained in Harris Hematoxylin tissues were cleared in Clarifier-2 and incubated with Bluing reagent (all from Richard-Allan Scientific). Slides were dehydrated in 70% and 100% ethanol and xylenes and mounted with Cytoseal 60. Hematoxylin-eosin staining was performed similarly by a standard method using Harris Hematoxylin followed by staining with eosin-phloxine solution as described before^26^. All slides were processed together to reduce batch effects.

### Image acquisition and analysis

Images for histological scoring were acquired with a Nuance Multispectral Imaging System (PerkinElmer) at 40x magnification using a spectral library created with slides with DAB or hematoxylin only staining. Six representative images of each specimen were captured. Ten percent of these images were used to create algorithms with inForm software v2.1.1. (PerkinElmer) to segment tissue area and nuclei. Cytoplasm was determined as a 20-pixel thick area around the nucleus. Four intensity intervals were manually established and used for categorical scoring of cytoplasmic staining (Supplemental Figure 1.). H-score was calculated by the software using the following formula: H-score=3x(% of 3+ cells)+2x(% of 2+ cells)+1x(% of 1+ cells). Representative images were taken with a Nikon Eclipse E600 microscope at 10x, and 40x magnification.

### Cell culture

Prostate stromal (BHPrS-1) and epithelial (NHPrE-1 and BHPrE-1) cells were gifts from Dr. Simon Hayward (NorthShore Research Institute, Evanston, IL). NHPrE-1 and BHPrE-1 lines were generated from primary cells by spontaneous immortalization, whereas BHPrS-1 were immortalized using hTERT^27^. THP-1 leukemia, LNCaP, and 22RV1 prostate cancer lines were purchased from ATCC. C4-2B^28^ cells were provided by Drs. Ruoxiang Wu and Leland Chung (Cedars-Sinai, Los Angeles, CA). NHPrE-1 and BHPrE-1 cells were cultured in DMEM:F12 in 1:1 ratio supplemented with 0.4% bovine pituitary extract, insulin-transferrin-selenium mix, epidermal growth factor (10ng/ml), antibiotic-antimycotic mix and 5% FBS (Thermo Fisher Scientific). THP-1, LNCaP, C4-2b and 22Rv1 cells were cultured in RPMI containing 10% FBS and BHPrS-1 cells with 5% FBS.

### Western Blot

To analyze the protein expression of OPN across cell types, cells were seeded in 6-well plates at a density of 100,000 cells/well and grown for 2 days. THP-1 cells were differentiated into M0 macrophages by adding 60 ng/ml phorbol myristate acetate to the growth medium for 2 days followed by resting in normal growth medium for another day^29^. Cells were washed with ice cold PBS 2 times and then 100 µl RIPA buffer (120mM NaCl, 50 mM pH 8.0 Tris, 0.5% NP-40, 1 mM EGTA) containing protease and phosphatase inhibitors (100 µg/mL PMSF, 1 mM NaOrVa, 50 µg/mL aprotinin, 50µg/mL leupeptin, all from Fisher Scientific) was added to the cells for 10-15 min. Cells were collected by scraping and cell debris was removed by centrifugation at 12,000 rpm at 4°C for 8 min.

Human prostate tissues (1-2 mm^3^) were transferred into 100 μl RIPA buffer/mm^3^ tissue, incubated for 1 h, homogenized with a pellet pestle and incubated for 1 h while kept on ice at all times. Debris was removed by centrifugation at 12,000 rpm for 8 min and supernatant was collected.

Protein concentration was determined using the Bio-Rad Protein Assay according to the manufacturer’s protocol with absorbance read at 595 nM on a SpectraMax ID3 spectrophotometer (Molecular Devices). Proteins were denatured in NuPage sample buffer containing 2.5 % β-mercaptoethanol and equal amounts (10-20 µg) were loaded onto 4-12% SDS-PAGE gels (NuPAGE). Proteins were transferred onto PVDF (Bio-Rad) membrane in 1X transfer buffer (NuPage Novex; NP-0006). Protein loading and transfer was confirmed with PonceauS staining (Boston BioProducts). Membranes were blocked with 10% milk in TBST for 1h at room temperature. Primary antibodies OPN (1:1000 dilution, ab8448, Abcam), anti-tubulin (1:2000 dilution, ab7291, Abcam) or anti-GAPDH (1:2000 dilution, AM4300, Ambion) were added overnight in 2.5% milk-fat. Secondary antibodies (Amersham) were used at 1:5000 dilution in 2.5% milk-fat for 1 hour. Bands were developed using the SuperSignal West Pico Plus Chemiluminescent Substrate and imaged with Bio-Rad Chemi-Doc Touch, then processed using Bio-Rad Image Lab software (ver6.0.1).

### Reverse transcription (RT) and quantitative polymerase chain reaction (qPCR)

Cells were seeded in 6-well plates at 100.000 cells/well density. The next day, recombinant human OPN (rhOPN) was added at 500 ng/ml concentration for 2, 4, 6 or 8 h or for 2 days (refreshing treatment on the second day) in RPMI containing 1 % heat-inactivated FBS for BHPrS-1 cells. For BHPrE-1 and NHPrE-1, cells were starved in serum-free media on the second day of culture and treatment of rhOPN was added on the third day for 2, 4, 6 or 8 h. RNA was isolated with the RNeasy minikit (Qiagen) and 1 µg of RNA was transcribed into cDNA using the iScript Kit (BioRad). The cDNA was then diluted 10x with water and 2 µl was mixed with iQ SYBR Green Supermix (BioRad) containing 100 nM of forward and reverse primers. The qPCR was performed on a CFX96 Touch (BioRad). Primer sequences are listed in Supplemental Table 2. Fold changes were calculated by the 2^-ΔΔCt^ method, normalized to untreated controls that were harvested at the same time point as OPN-treated cells (2,4,8 and 12 h for Figure 5, Table 1. and Supplemental Table 1. and 2d for Figure 6 and Table 2) and using *GAPDH* as a housekeeping gene.

**Figure 5.**
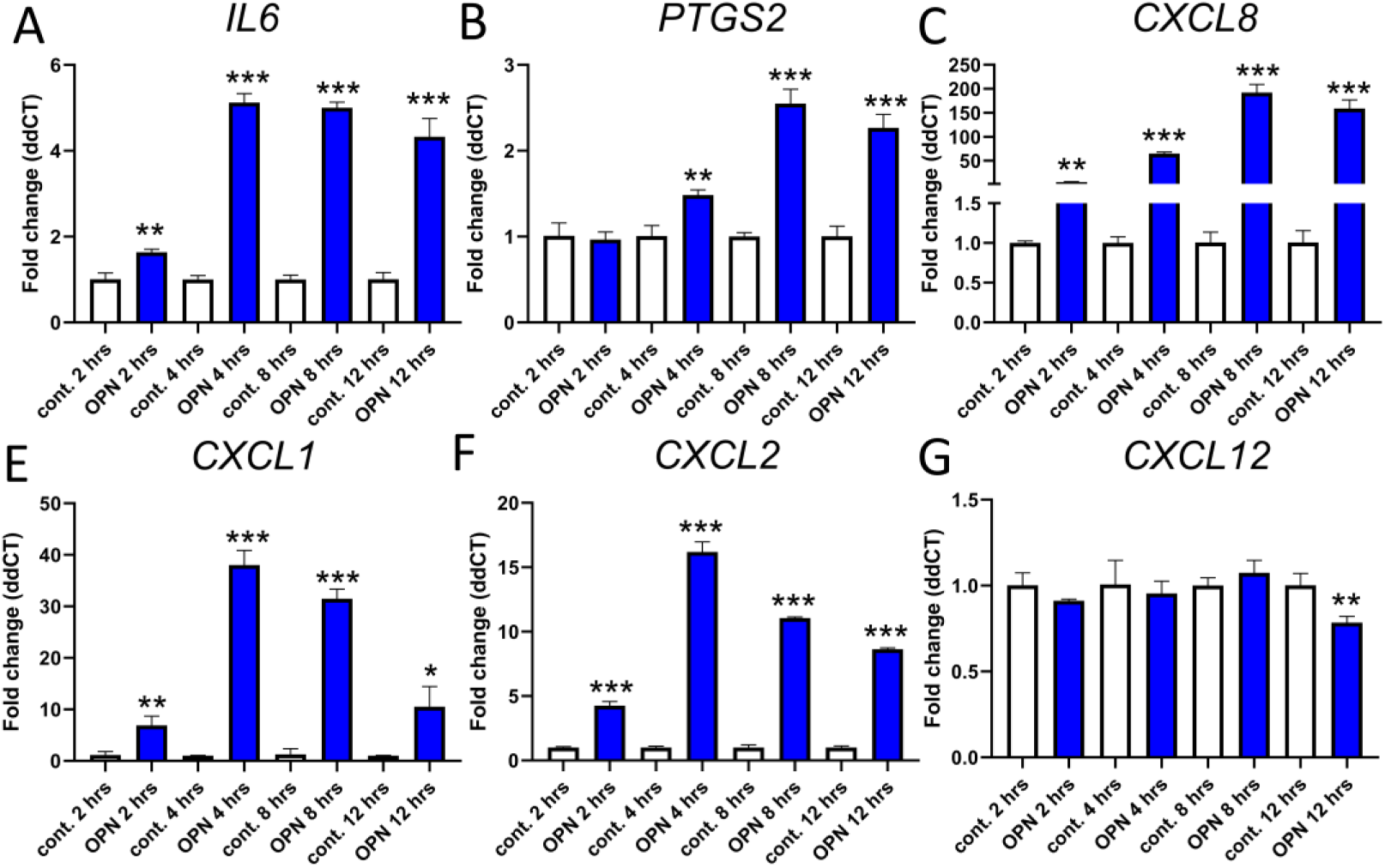
OPN induces the expression of various inflammatory genes in BHPrS-1 stromal cells. OPN induced the expression of *IL-6* (**A**)*, PTGS2* (**B**)*, CXCL8* (**C**)*, CXCL1* (**D**)*, CXCL2* (**E**) but not *CXCL12* (**F**). Expressional changes were detected at 2, 4, 8, or 12 hours after the addition of 500 ng/ml rhOPN whereas controls were generated for each time point incubated with the treatment medium. Results are representative of 2 independent experiments. Mann-Whitney non-parametric test was employed to determine significance versus time-matched controls. *p<0.05, **p<0.01, ***p<0.001.

**Figure 6.**
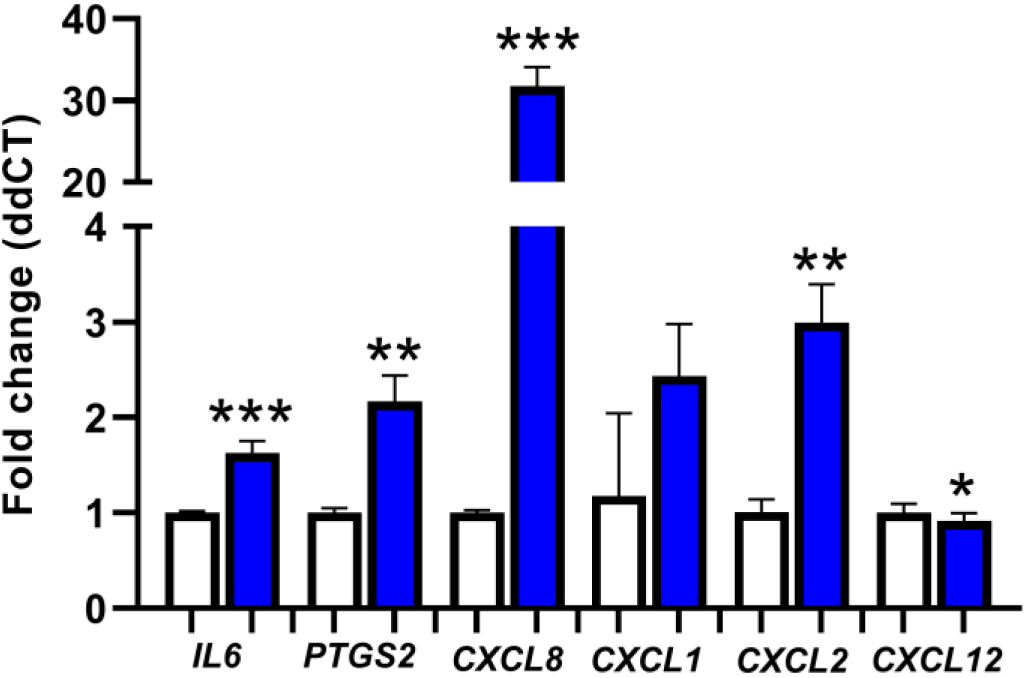
The cytokine transcriptional pattern is preserved with extended OPN treatment. *IL-6, PTGS2, CXCL8,* and *CXCL2* gene expression levels remained significantly elevated after two consecutive 24-hour treatments with 500 ng/ml OPN. *CXL12* levels decreased similarly to the short-term treatments. Mann-Whitney non-parametric test was employed to determine significance versus vehicle treated controls. *p<0.05, **p<0.01, ***p<0.001.

**Table 1.** Fold-changes in genes examined in response to short-term treatments with rhOPN in BHPrS-1 cells.

| gene | fold elevation (ddCT) |  |  |  |  |  |  |  |
| --- | --- | --- | --- | --- | --- | --- | --- | --- |
|  | cont. | OPN 2 hrs | cont. | OPN 4 hrs | cont. | OPN 8 hrs | cont. | OPN 12 hrs |
| *IL6* | 1.01±0.15 | **1.64±0.07\*\*** | 1.00±0.09 | **5.12±0.21\*\*\*** | 1.00±0.10 | **5.00±0.13\*\*\*** | 1.01±0.15 | **4.33±0.43\*\*\*** |
| *PTGS2* | 1.01±0.15 | 0.97±0.09 | 1.01±0.12 | **1.48±0.06\*\*** | 1.00±0.05 | **2.55±0.17\*\*\*** | 1.00±0.12 | **2.27±0.16\*\*\*** |
| *CXCL8* | 1.00±0.02 | **5.26±1.13\*\*** | 1.00±0.08 | **64.94±3.06\*\*\*** | 1.01±0.13 | **192.08±17.01\*\*\*** | 1.01±0.15 | **158.8±17.87\*\*\*** |
| *CXCL1* | 1.13±0.70 | **6.91±1.79\*\*** | 1.00±0.07 | **38.04±2.83\*\*\*** | 1.28±1.07 | **31.49±1.87\*\*\*** | 1.00±0.05 | **10.54±3.91\*** |
| *CXCL2* | 1.00±0.07 | **4.25±0.33\*\*\*** | 1.00±0.11 | **16.20±0.78\*\*\*** | 1.01±0.19 | **11.06±0.08\*\*\*** | 1.01±0.13 | **8.64±0.10\*\*\*** |
| *CXCL12* | 1.00±0.07 | 0.91±0.01 | 1.01±0.14 | 0.95±0.07 | 1.00±0.05 | 1.07±0.07 | 1.00±0.07 | **0.78±0.04\*\*** |
| *ACTA2* | 1.00±0.07 | 1.1±0.06 | 1.00±0.07 | 1.10±0.07 | 1.00±0.12 | 1.05±0.06 | 1.00±0.09 | 0.95±0.02 |
| *SPP1* | 1.00±0.05 | 0.80±0.15 | 1.01±0.21 | 1.38±0.23 | 1.01±0.18 | 1.08±0.23 | 1.01±0.14 | 0.89±0.12 |
| *TGFB1* | 1.00±0.04 | 1.07±0.02 | 1.00±0.07 | 0.99±0.05 | 1.00±0.08 | 1.10±0.06 | 1.00±0.07 | 0.89±0.19 |
| *MMP1* | 1.01±0.17 | 0.97±0.12 | 1.00±0.08 | 1.04±0.02 | 1.00±0.05 | 1.14±0.20 | 1.00±0.11 | 1.07±0.18 |
| *MMP2* | 1.00±0.03 | 1.02±0.05 | 1.00±0.12 | 1.01±0.05 | 1.00±0.08 | 1.15±0.10 | 1.00±0.09 | 0.84±0.10 |
| *MMP9* | 1.00±0.04 | **1.16±0.01\*\*** | 1.01±0.14 | 1.19±0.10 | 1.00±0.07 | 1.36±0.21 | 1.01±0.16 | 1.26±0.08 |
| *TIMP1* | 1.00±0.04 | 1.07±0.02 | 1.00±0.07 | 0.99±0.05 | 1.00±0.08 | 1.10±0.06 | 1.00±0.07 | 0.89±0.19 |
| *COL1A1* | 1.01±0.14 | 0.75±0.05 | 1.02±0.22 | 0.84±0.05 | 1.00±0.06 | 1.17±0.10 | 1.00±0.04 | 0.96±0.11 |
| *Col1A2* | 1.00±0.08 | 1.05±0.04 | 1.00±0.11 | 1.00±0.06 | 1.00±0.05 | 1.10±0.18 | 1.00±0.05 | 0.89±0.07 |

Fold-elevation values are shown ± SD, 2, 4, 8, or 12 hours after the addition of 500 ng/ml rhOPN. A distinct control was generated for each time point using treatment medium only. Results are representative of 2 independent experiments. Mann-Whitney non-parametric test was employed to determine significance. \*\*p<0.01, \*\*\*p<0.001.

**Table 2.** The effects of long-term (2 day) OPN treatment on gene expression.

| fold elevation (ddCT) |  |  |  |  |  |  |  |  |
| --- | --- | --- | --- | --- | --- | --- | --- | --- |
| gene | cont. | OPN | gene | cont. | OPN | gene | cont. | OPN |
| *IL6* | 1.00±0.02 | **1.63±0.12\*\*\*** | *CXCL12* | 1.00±0.09 | **0.92±0.08\*** | *MMP2* | 1.00±0.07 | 1.09±0.04 |
| *PTGS2* | 1.00±0.05 | **2.17±0.27\*\*** | *ACTA2* | 1.00±0.04 | **1.29±0.16\*** | *MMP9* | 1.01±0.16 | 1.08±0.13 |
| *CXCL8* | 1.00±0.03 | **31.76±2.31\*\*\*** | *SPP1* | 1.06±0.40 | 1.04±0.21 | *TIMP1* | 1.00±0.07 | 1.13±0.13 |
| *CXCL1* | 1.18±0.86 | 2.43±0.55 | *TGFB1* | 1.00±0.08 | 1.02±0.10 | *COL1A1* | 1.01±0.15 | 1.08±0.01 |
| *CXCL2* | 1.01±0.14 | **2.99±0.40\*\*** | *MMP1* | 1.00±0.12 | **1.73±0.09\*\*\*** | *Col1A2* | 1.00±0.06 | 1.10±0.02 |

Fold-elevation values are shown ± SD, following two 24-hour treatments with 500 ng/ml rhOPN. Results are representative of two independent experiments. Mann-Whitney non-parametric test was employed to determine significance. \*p<0.05, \*\*p<0.01, \*\*\*p<0.001.

### Enzyme-linked immunosorbent assay (ELISA)

BHPrS-1 or NHPrE-1 cells were seeded in 6-well plates at 100,000 cells/well density. Treatments IL-1β (1, 10 or 100 pg) or TGF-β1 (0.1, 1 or 10 ng) were added in RPMI supplemented with 1% heat-inactivated FBS for BHPrS-1 and DMEM/F12 with 0.1% BSA for NHPrE-1 cells. Media were collected 24 hours later and cell debris was removed by centrifugation at 3000 rpm for 3 min and stored at −80°C. The concentration of OPN was determined by the Human Osteopontin (OPN) DuoSet ELISA (DY1433, R&D Systems) according to the manufacturer’s instructions. OPN concentration was calculated by GraphPad Prism ver. 8 using second order polynomial regression analysis.

### Statistical analysis

Statistical analysis was performed with GraphPad Prism using non-parametric Kruskal-Wallis and Mann-Whitney tests. Results were considered significant at p<0.05. All experiments were repeated at least once.

## Results

### OPN levels are increased in advanced BPH

We quantified OPN immunostaining in two patient populations, referred to as incidental and surgical BPH (I-BPH and S-BPH), to determine whether prostatic OPN abundance associates with symptomatic disease. Surgical BPH (S-BPH) specimens were prostate transition zone specimens from surgeries to relieve symptoms or obstruction due to LUTS. Some S-BPH patients received 5-α reductase inhibitors (5ARI, n=6), α-blockers (a-B, n=10), a combination of 5ARI and a-B (5ARI+a-B, n=7) or no treatment (n=7) for BPH/LUTS. I-BPH specimens were prostate transition zone specimens from mildly symptomatic patients undergoing radical prostatectomies for peripheral zone-confined prostate cancer. S-BPH was previously demonstrated to show signs of BPH progression such as loss of smooth muscle differentiation and fibrosis compared to I-BPH^25^.

OPN expression was visualized by immunohistochemistry (Figure 1) and staining was quantified by H-scoring. We observed significantly more OPN in S-BPH than in I-BPH (p=0.0107, Mann-Whitney test, Figure 2A). We then further divided S-BPH specimens by treatment (no treatment, 5ARI, a-B, or 5ARI+a-B). OPN immunostaining was most abundant in the 5ARI+a-B group and was significantly higher than I-BPH (p=0.0289, Mann-Whitney test, Figure 2B, Supplemental Figure 2.), however, the difference between S-BPH treatment groups was not significant.

**Figure 1:**
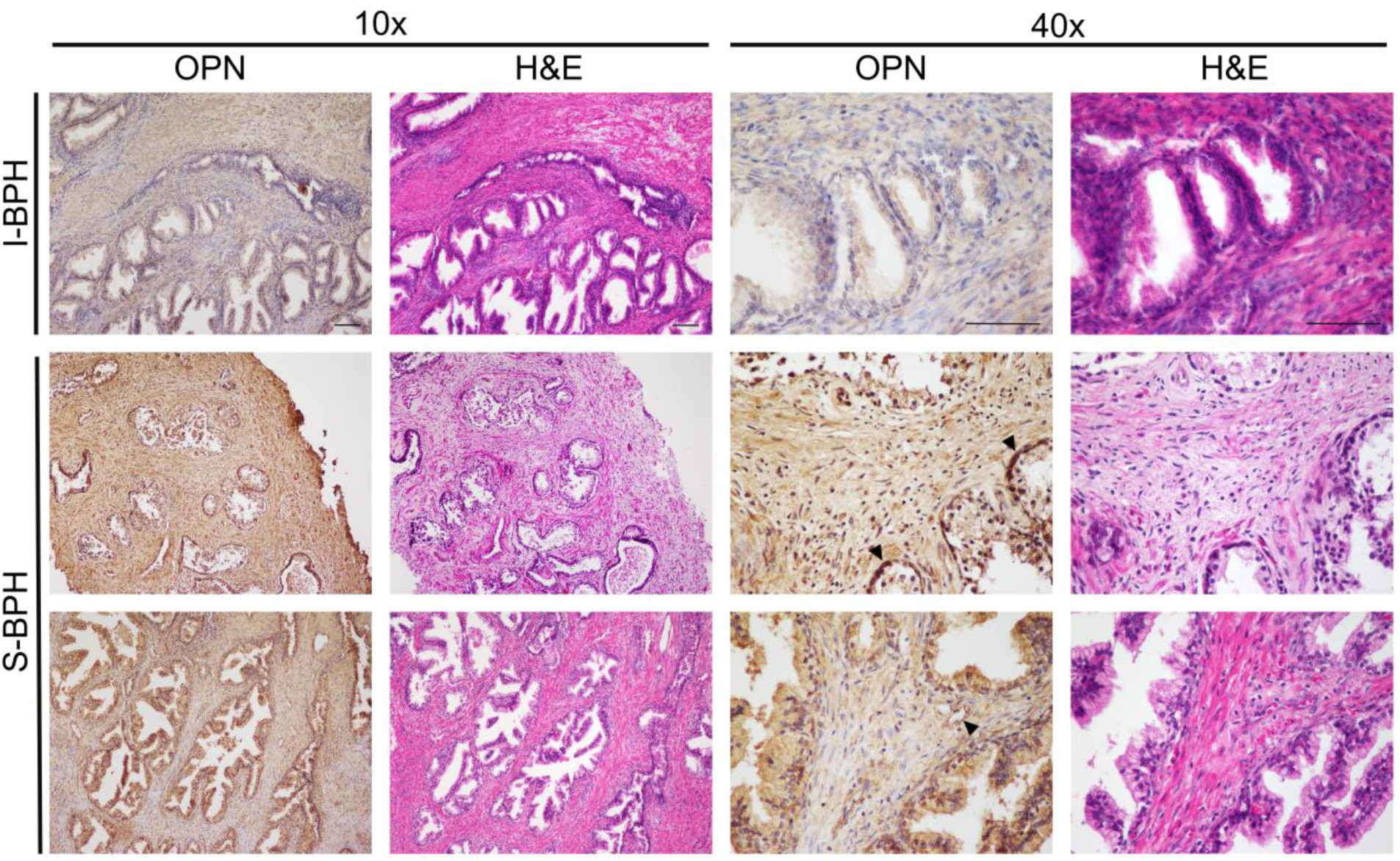
OPN protein levels are localized to multiple cell types in BPH specimens. Representative images of OPN IHC performed on incidental and surgical BPH (I-BPH and S-BPH) specimens are shown at 10x, and 40x magnification. OPN has the highest levels in epithelial cells with an occasional upregulation in basal cells (indicated by arrows) and in endothelial cells (asterisk). It is also distributed throughout the stroma with varying density.

**Figure 2:**
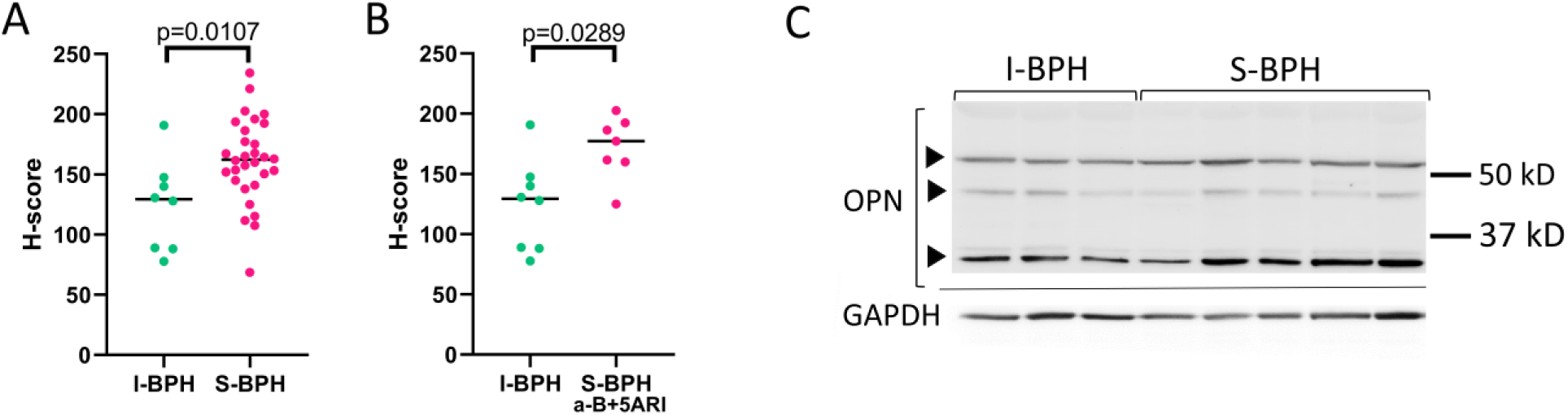
OPN protein expression is significantly increased in symptomatic BPH. (**A**) Determination of H-score using inForm revealed a significant increase in OPN protein expression in surgical (S-BPH, n=24) compared to incidental BPH (I-BPH, n=8, p=0.0107). (**B**) OPN levels were highest in patients receiving the combination of α-blockers (a-B) and 5α-dehydrogenase inhibitors (5ARI) compared to I-BPH (p=0.0289). The complete subgroup analysis containing all treatment groups treatments (a-B;, 5ARI; their combination or no treatment) are shown in Supplemental Figure 2. Mann-Whitney or Kruskall-Wallis non-parametric tests were employed to determine significance. Horizontal lines on graphs indicate median. (**C**) Western blot analysis identified 3 different variants of OPN, including the cleaved form at 32 kD which showed the most pronounced elevation in S-BPH. GAPDH was used as a loading control.

Representative IHC images reveal (Figure 1) OPN immunostaining in glandular cells, with occasional stronger intensity in basal cells, diffusely or as deposits in the stromal compartment, in endothelial cells and in some inflammatory cells. We used Western blotting as a secondary method to detect OPN protein abundance and found higher levels in S-BPH compared to I-BPH (Figure 2C). The polyclonal antibody used in our studies (ab8448, Abcam) was raised against a synthetic peptide corresponding to the C-terminal of OPN protein that is present in all splice variants and in OPN fragments generated by thrombin, MMP-3 and MMP-9 cleavage^30^. Two intact OPN peptides (42 and 52 kD) were detected but the identification of a specific splice variant is challenging due to the high level of posttranslational modifications. We also detected high levels of cleaved OPN at 32 kD.

### OPN is expressed in prostate cells and its secretion is stimulated by inflammatory signals

Most studies involving OPN synthesis in the prostate have focused on cancer cells and macrophages^31–35^. Therefore, our aim was to determine whether non-transformed resident prostate cells also synthesize OPN. We first tested whether OPN protein is expressed by benign prostate stromal (BHPrS-1), epithelial (NHPrE-1 and BHPrE-1) and cancer cell lines (LNCaP, C4-2B and 22RV1) (Figure 3A). Epithelial cell lines were grown as monolayers or were embedded in matrigel to differentiate and form organoids^36^. We detected three intact OPN peptides (42, 52 and 70 kD) in all cells, as well as the 32 kD cleaved peptide in BHPrS-1 cells.

**Figure 3:**
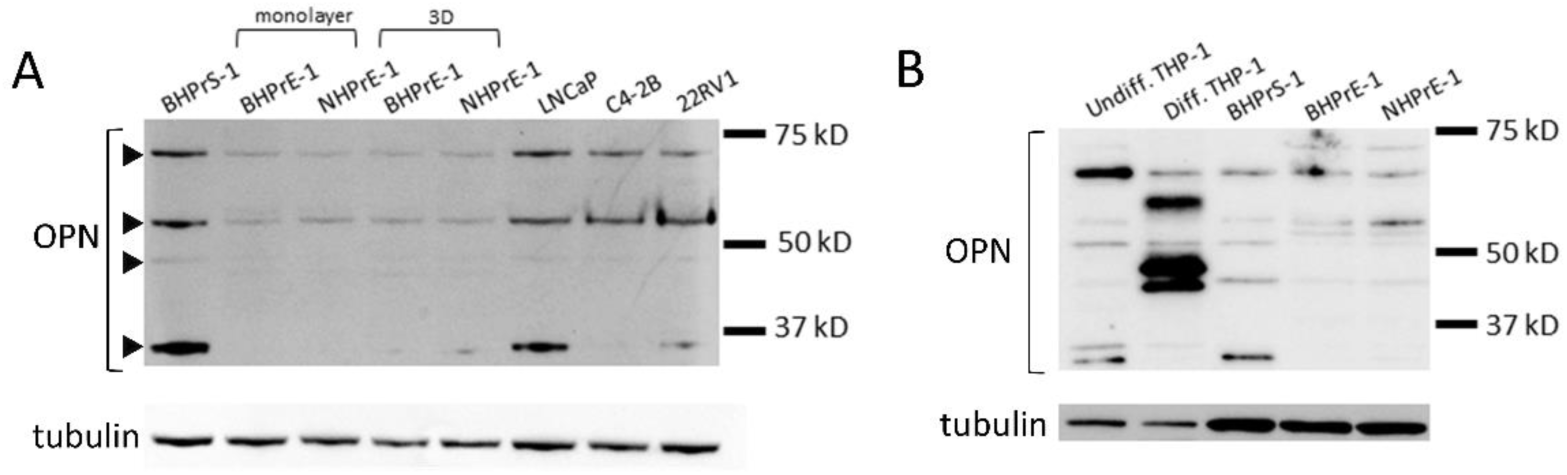
OPN is expressed in benign prostate epithelial and stromal cell lines. OPN expression was detected by Western blot in lysates of immortalized prostate stromal (BHPrS-1) and epithelial cell lines (BHPrE-1 and NHPrE-1, monolayer or 3D matrigel cultures grown for 8 days). (**A**) Benign cells express similar OPN variants as prostate cancer cell lines LNCaP, C4-2B and 22RV1, however, at lower levels and lacking the cleaved variant at 32 kD. (**B**) Undifferentiated THP-1 leukemia cells express OPN variants most similar to BHPrS-1 cells, however, upon their differentiation by phorbol myristate acetate to M0, unique OPN forms appear.

Macrophages express various OPN splice variants including the intracellular variant produced by an alternative translation initiation site^30, 37^. To compare OPN variants expressed in prostate cells to those produced by macrophages, we evaluated OPN variants in THP-1 cells monocytes, before and after converting them into M0 macrophages with phorbol 12-myristate 13-acetate^29^. OPN variants in undifferentiated THP-1 cells were the same as those detected in prostate stromal and epithelial cells (Figure 3B). In contrast, in converted M0 macrophages, we observed a switch to unique OPN variants at 48 and 68 kD as well as an increase in the 42 kD product. We conclude that relative abundance of OPN variants changes during macrophage activation. Whether this change is caused by alterations in splice variant transcription or posttranscriptional modifications remains to be determined.

We next tested whether OPN is secreted by prostate cells. Basal OPN secretion, as determined by ELISA in media after 24 hours of incubation, was approximately 5-fold higher in NHPrE-1 epithelial than BHPrS-1 stromal cells (Figure 4). BHPrS-1 cell secretion of OPN was increased 2-fold by IL-1β and 2.5-fold by TGF-β1. In contrast, NHPrE-1 cell secretion of OPN was increased by TGF-β1 but not IL-1β. These results support the concept that prostatic stromal and epithelial cells can autonomously secrete OPN in response to pro-inflammatory stimuli.

**Figure 4.**
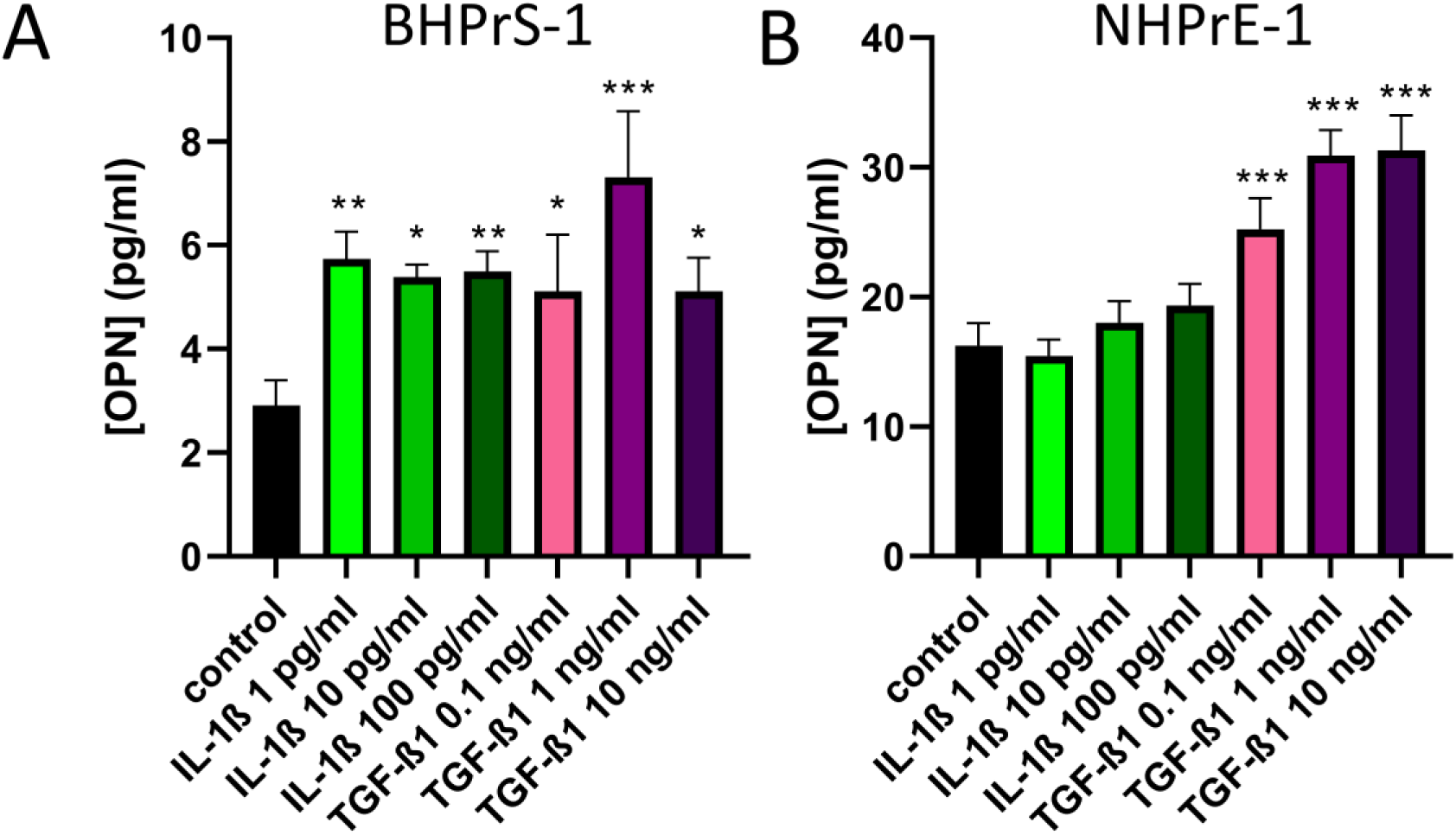
Prostate cellssecrete OPN in response to cytokines IL-1β and TGF-β1. (**A**) BHPrS-1 cells have increased OPN secretion in response to both IL-1β and TGF-β1 whereas (**B**) in NHPrE-1 cells, only TGF-β1 stimulated the secretion of OPN. Results are representative of at least 2 independent experiments. Kruskall-Wallis non-parametric test was employed to determine significance versus control. *p<0.05, **p<0.01, ***p<0.001.

### OPN stimulates cytokine production in prostate stromal cells

We next analyzed the expression of previously described or hypothetical downstream targets of OPN in benign prostate cells by qPCR. Initial experiments identified a 500 ng/ml OPN concentration as optimal (versus 100 ng/ml, 500 ng/ml, 1 µg/ml) for inducing OPN target gene expression (results not shown). BHPrS-1, BHPrE-1 and NHPrE-1 cells were incubated with 500 ng/ml recombinant human (rh)OPNfor 2, 4, 8 and 12 hours. We examined the abundance of OPN target genes linked to tissue remodeling and fibrosis in pulmonary fibrosis, muscular dystrophy, aortic injury and cancer^24, 38–40^ including *MMP1, MMP2, MMP9, TIMP1, COL1A1, COL1A2, TGFB1, and ACTA2*. Surprisingly, no significant change was observed in the expression of these genes in response to OPN except a 16% increase in MMP9 mRNA expression after 2 hours of treatment in BHPrS-1 cells (Table 1). We also tested whether the expression of*TGFB1*, *MMP1* and *TIMP1* was affected in BHPrE-1 and NHPrE-1 cells after OPN treatment but no significant change was detected (Supplemental Table 1).

We surmised that OPN might influence the abundance of a unique set of target genes in BPH, including those already linked to BPH pathogenesis such as *IL6*, *CXCL8*, *CXCL1*, *CXCL2*, *CXCL12*^41–43^. We examined the impact of OPN exposure on target gene expression in BPH stromal-derived BHPrS-1 cells. The mRNA expression of all gene listed above except *CXCL12* was significantly increased 2 h after initiation of OPN treatment and continued to rise through 8 h of OPN treatment (Figure 5). OPN elicited the greatest magnitude of change for *CXCL8* mRNA, which was increased to 192-fold elevation (p<0.05) 8 h after the initiation of OPN treatment. We also investigated the mRNA expression of prostaglandin-endoperoxidase synthase 2 (*PTGS2*), an inflammation biomarker, induced by numerous inflammatory signals^44, 45^. *PTGS2* mRNA expression increased 4 h after initiation of OPN treatment (1.48-fold, p<0.01) and peaked at 8 h (2.55-fold, p<0.001). Interestingly, *CXCL12* mRNA expression was slightly but significantly reduced 12 h after OPN treatment (0.78-fold, p<0.01).

We next tested whether OPN drives *PTGS2* expression in BHPrE-1 and NHPrE-1 cells. OPN increased *PTGS2* mRNA expression (1.15-fold, p<0.01), 8 h after initiation of treatment in BHPrE-1 cells, but not in NHPrE-1 cells and at no other time points (Supplemental Table 1.). We then tested whether OPN increased mRNA expression of three other chemokines (*CXCL8*, *CXCL1*, *CXCL2*) but only slight elevations were observed (Supplemental Table 1). We conclude that the primary OPN-initiated inflammatory response is mostly mediated through prostate stromal cells.

We next tested long-term responses to OPN exposure. BHPrS-1 cells were exposed to rhOPN for 2 d (Figure 6 and Table 2). *IL6*, *PTGS2*, *CXCL8*, and *CXCL2* mRNA expression remained significantly elevated (1.63-, 2.17-, 31.76-, 2.99-fold, respectively) and *CXCL12* expression was decreased (0.92-fold, p<0.05) compared to untreated controls. Interestingly, levels of *ACTA2* and *MMP1* mRNA were significantly increased by OPN (1.29-fold and 1.73-fold, respectively); whereas the mRNA expression of *TGFB1*, *SPP1*, *MMP2*, *MMP9*, *TIMP1*, *COL1A1* and *COL1A2* mRNAs was unchanged by OPN (Table 2).

## Discussion

We provide the first direct evidence that OPN is more abundant in S-BPH than I-BPH specimens, suggesting a potential role for OPN in the development of BPH and associated voiding sequelae. As reported by previous investigations^25, 46^, S-BPH shows signs of BPH progression including loss of smooth muscle differentiation and higher levels of fibrosis compared to the prostates of mildly symptomatic I-BPH patients, implying that OPN may have an important role in BPH progression. Previous studies of OPN in prostatic diseases primarily focused on malignancies and often used BPH prostate as a control. These studies reported weak staining/expression of OPN in BPH specimens compared to those with malignant disease^31, 32^. However, other studies have reported the upregulation of OPN in BPH versus normal prostate or healthy controls. Thalmann et al. demonstrated a comparable OPN mRNA expression level in prostate cancer and BPH^33^. Furthermore, serum OPN levels were found to be significantly elevated in BPH patients compared to age-matched healthy individuals with no history of prostate or metabolic bone disease^34^. In addition, OPN immune-reactivity as detected by the occurrence of serum OPN antibodies thus indicating a development of autoimmunity against OPN, is more prevalent in BPH patients compared to healthy donors^47^. According to these findings, there is an association between OPN in benign prostatic diseases; however, how OPN could be contributing to BPH was not evaluated.

Consequently, our initial goal was to clarify whether prostatic OPN expression correlates with the progression of BPH. For this, we compared OPN expression as determined by H-score in patients with advanced BPH who have undergone surgery to relieve LUTS to the transition zone of low-grade, low-volume peripheral zone-restricted prostate cancer in patients^46, 48^. We found higher OPN levels in S-BPH specimens. We also found when patients were stratified based on medical treatments, prostatic OPN levels were highest in patients who received an a-B and 5ARI combined treatment. Remarkably, patients who progress to surgery failing the combination of a-B and 5ARI were shown to have higher prostatic levels of collagens^15^, which supports the concept that mediators of fibrosis, such as OPN, are important in BPH-progression.

In our study, the predominant variant of OPN observed in human tissues and stromal cells was a cleaved product. Such OPN fragments often attain higher activity; cleavage by thrombin exposes a cryptic integrin-binding motif^49^ which leads to increased activation, proliferation, and migration of hepatic stellate cells leading to increased fibrogenesis^50^. This variant was not investigated by previous studies focusing on prostatic diseases.

Fascinatingly, intracellular protein levels of OPN in BHPrS-1 stromal cells were comparable to prostate cancer cell lines LNCaP, 22RV1 and C4-2B whereas BHPrE-1 and NHPrE-1 benign epithelial cells possessed visibly lower levels. This is likely due to an increased secretion level in epithelial cells, which is supported by our ELISA measurements showing an approximately 3-fold higher basal level of OPN in medium produced by NHPrE-1 than BHPrS-1 cells. In addition, while a previous study deciphering the role of OPN in the proliferation of epithelial cells identified OPN as a macrophage-released cytokine^35^, our study showed that resident prostate cells are also able to secrete OPN in a process augmented by cytokines (IL-1β and TGF-β1).

The role of prostate stromal cells in regulating inflammatory processes is an intriguing area of research. Stromal cells express toll-like receptors responsible for recognizing microbes and are also demonstrated to act as antigen-presenting cells^41^. They express costimulatory factors and receptors required for the activation of CD4^+^ T-cells including IL-12, CD40, CD80, CD86 and CD134^41^. Interestingly, our study showed that OPN stimulates the expression of *IL6* and *IL8*, which potentially leads to increased secretion of these cytokines similarly to the action of IFNγ and IL-17^41^. Overexpression of IL-8 in prostatic epithelial cells leads to hyperplasia and the development of periglandular reactive stroma with increased pro-collagen-1 and tenascin levels consistent with fibrosis^51^. In addition, OPN also stimulated the expression of *CXCL1* and *CXCL2*, but not *CXCL12*. Prostate stromal cells have been previously shown to secrete CXCL1 and CXCL2 in response to IL-1β^52^. Moreover, the activation of CXCL1, CXCL2 and CXCL8 receptors CXCR1 and CXCR2^53^ have been shown to induce the expression of COL1A1 and ACTA2^54^. CXCR1 and CXCR2 are expressed on neutrophils, T-cells and mast cells^55, 56^ implying that these cells are all potentially attracted by prostate stromal cell-derived chemokines in response to increased levels of OPN. It also has to be acknowledged, that although there was a slight elevation in the expression of *ACTA2*, a marker for myofibroblast phenoconversion, *COL1A1* levels did not follow the same pattern and it is likely that the fibrotic effect of OPN may require the accumulation and extended presence of its target cytokine genes. Nevertheless, these findings indicate that OPN may initiate proinflammatory pathways and attract immune cells to the prostate leading to chronic inflammation and fibrogenesis.

The lack or limited induction of genes directly involved in fibrosis, including *COL1A1*, *COL1A2*, *TGFB1*, and tissue remodeling, including MMPs, in our studies were contradictory to previous findings in fibrotic models^38, 39, 57^. However, a great portion of studies linking OPN to collagen production and fibrosis utilized the OPN-KO mouse as a model^22, 57, 58^. In bleomycin-induced dermal fibrosis, OPN-KO mice had reduced number of multiple cell types, including myofibroblasts, decreased collagen content and inflammation. Interestingly, in this study, the TGF-β pathway activation was only affected *in vivo* but this functional disturbance was not reproducible *in vitro* ^57^. This may suggest that the fibrotic action of OPN may only manifest *in vivo* as a cumulative product of stromal, epithelial and immune cell interaction s in the prostate. Indeed, our data suggests a paracrine feedback mechanism where primarily epithelial cells secrete OPN and stromal cells respond by up-regulating pro-inflammatory cytokines, in turn inducing more OPN secretion by epithelial cells (Figure 7). Even though our pathological evaluation determined no significant difference in inflammation between I-BPH and S-BPH, a former study identified an increased activation of the inflammatory response transcriptional network in S-BPH using a sensitive microarray method^25^. Abundance of inflammatory infiltrates in the prostate also correlates with symptom score and prostate volume^59^. Our proposed mechanism for OPN in the exacerbation of inflammatory signals in the prostate may have a pivotal role in the progression of LUTS, however, more research is needed to reveal the exact contribution of OPN.

**Figure 7.**
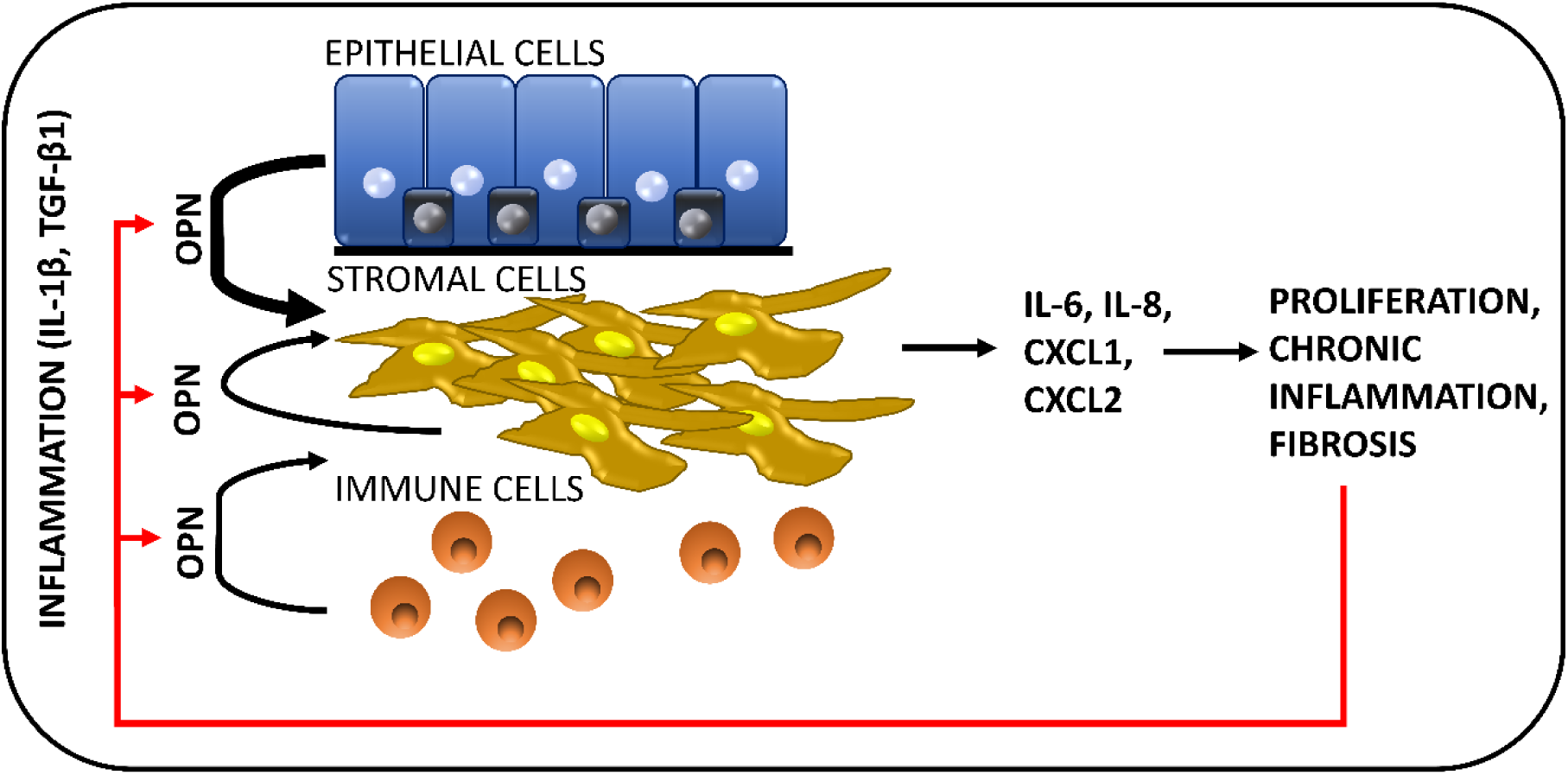
OPN exacerbates inflammation and fibrosis in the prostate. OPN secretion is stimulated by inflammatory cytokines including IL-1β and TGF-β primarily from epithelial cells. OPN stimulates the proliferation of epithelial cells and further promotes inflammation by triggering cytokine expression in stromal cells. Sustained inflammation will lead to fibrosis and the potentiation of the release of further OPN.

Lastly, we did not find marked responses (less than 30% elevation) in the expression of genes selected in this study in epithelial cells. However, OPN has been found to induce the proliferation of prostate epithelial cells via activation of Akt and ERK phosphorylation in immortalized prostate epithelial cells^35^ and it is likely that the proinflammatory action of OPN is primarily linked to stromal cells.

## Conclusions

Our study identified that OPN protein levels are higher in S-BPH, representing a more progressed, symptomatic stage of the disease, compared to mildly symptomatic I-BPH. We show that OPN is expressed across diverse prostate cell types including immune, epithelial, stromal and endothelial cells. We demonstrate that prostate epithelial and stromal cells can autonomously synthesize OPN in response to proinflammatory stimuli. We also show that OPN drives the expression of multiple cytokines and chemokines within prostate stromal cells, linking it directly to the proinflammatory and pro-fibrotic pathways that were previously associated with male LUTS and suggesting that OPN can exacerbate inflammatory processes in the prostate. Therapeutics targeting OPN action may have multiple beneficial activities in BPH, including anti-inflammatory, anti-fibrotic and anti-proliferative effects; however, more research is needed to identify the molecular mechanism by which OPN contributes to BPH.

## Acknowledgements

BPH tissue was provided by the NCI funded Cooperative Human Tissue Network. Use of human tissue is approved by the VUMC (#120944), the CWRU (#STUDY20190025) and the UW (2019-0912) IRBs. We thank the patients who have generously donated tissue for this study. The study was supported by a start-up fund provided by the Department of Urology, Case Western Reserve University (to MMG), NIH NIDDK K12 DK100022-06 (to PP), T32GM008056-37 (to SEK), 5R01 DK111554-03 (to RJM) and U54 DK104310 (to WAR).

