## Supplemental Information for "Prostatic osteopontin expression is associated with symptomatic benign prostatic hyperplasia"

| Supplemental Table 1. Benign epithelial cell lines show only mild expressional changes in response to OPN. | | | | | | | | | |
| --- | --- | --- | --- | --- | --- | --- | --- | --- | --- |
| NHPrE-1 |  | **fold elevation (ddCT)** | | | | | | | |
|  | **gene** | **cont.** | **OPN 2 hrs** | **cont.** | **OPN 4 hrs** | **cont.** | **OPN 8 hrs** | **cont.** | **OPN 12 hrs** |
|  | *PTGS2* | 1.00±0.09 | 1.09±0.10 | 1.01±0.15 | 1.22±0.25 | 1.01±0.15 | 1.08±0.12 | 1.00±0.05 | 1.13±0.23 |
|  | *CXCL8* | 1.00±0.05 | **1.14±0.05*** | 1.02±0.26 | 1.39±0.39 | 1.00±0.10 | 1.12±0.02 | 1.00±0.00 | 1.16±0.21 |
|  | *CXCL1* | 1.00±0.04 | 1.02±0.05 | 1.02±0.22 | 1.25±0.24 | 1.01±0.14 | 1.17±0.09 | 1.00±0.03 | 1.10±0.09 |
|  | *CXCL2* | 1.00±0.09 | 1.37±0.17 | 1.02±0.27 | 1.43±0.52 | 1.00±0.10 | 1.14±0.04 | 1.00±0.01 | 1.04±0.04 |
|  | *TGFB1* | 1.00±0.02 | 1.01±0.08 | 1.00±0.05 | 0.97±0.10 | 1.00±0.04 | 0.92±0.01 | 1.00±0.06 | 0.97±0.05 |
|  | *MMP1* | 1.00±0.07 | 0.92±0.07 | 1.00±0.05 | 0.99±0.06 | 1.00±0.11 | 1.10±0.01 | 1.00±0.04 | 0.98±0.06 |
|  | *TIMP1* | 1.00±0.03 | 1.03±0.05 | 1.00±0.12 | 1.01±0.03 | 1.00±0.05 | 1.02±0.03 | 1.00±0.01 | 1.05±0.05 |
| BHPrE-1 | *PTGS2* | 1.01±0.17 | 1.26±0.03 | 1.01±0.13 | 0.94±0.10 | 1.00±0.04 | **1.15±0.02**** | 1.00±0.06 | 1.07±0.06 |
|  | *CXCL8* | 1.00±0.07 | 1.19±0.10 | 1.00±0.11 | 1.11±0.21 | 1.00±0.02 | **1.05±0.01*** | 1.00±0.02 | 1.08±0.07 |
|  | *CXCL1* | 1.00±0.06 | 1.03±0.11 | 1.01±0.14 | 1.14±0.30 | 1.00±0.07 | 1.09±0.05 | 1.00±0.04 | 1.10±0.13 |
|  | *CXCL2* | 1.01±0.13 | **1.27±0.02*** | 1.00±0.10 | 1.28±0.39 | 1.00±0.04 | **1.20±0.03**** | 1.00±0.06 | 1.03±0.04 |
|  | *TGFB1* | 1.00±0.06 | 0.98±0.02 | 1.00±0.06 | 1.06±0.03 | 1.00±0.08 | 1.06±0.05 | 1.00±0.06 | 1.07±0.04 |
|  | *MMP1* | 1.00±0.03 | 1.08±0.08 | 1.00±0.10 | 1.00±0.10 | 1.00±0.06 | **1.10±0.01*** | 1.00±0.06 | 1.07±0.07 |
|  | *TIMP1* | 1.00±0.03 | 1.04±0.11 | 1.00±0.08 | 1.04±0.07 | 1.00±0.05 | 1.14±0.05 | 1.00±0.05 | **1.16±0.02**** |

Fold-elevation values are shown ± SD, 2, 4, 8, or 12 hours after the addition of 500 ng/ml rhOPN. A distinct control was generated for each time point using treatment medium only. Results are representative of two independent experiments. Mann-Whitney non-parametric test was employed to determine significance. *p<0.05, **p<0.01

| Supplemental Table 2. Primer sequences | | | | | |
| --- | --- | --- | --- | --- | --- |
| gene name | **Forward primer** | **Reverse primer** | **product length (bp)** | **Annealing temperature** | **reference** |
| Col1a1 | ACGAAGACATCCCACCAATC | ATGGTACCTGAGGCCGTTC | 50 | 58 | ^1^ |
| Col1a2 | GCATTCGTGGCGATAAGGG | CCATGGTGACCAGCGATACC | 109 | 58 |  |
| CXCL1 | CTGGCTTAGAACAAAGGGGCT | TAAAGGTAGCCCTTGTTTCCCC | 112 | 58 |  |
| CXCL12 | TGTGCCCTTCAGATTGTAGCC | ACTTTAGCTTCGGGTCAATGC | 70 | 58 |  |
| CXCL2 | GAAAGCTTGTCTCAACCCCG | TGGTCAGTTGGATTTGCCATTTT | 82 | 58 |  |
| CXCL8 | GAAGTTTTTGAAGAGGGCTGAGA | TTTGCTTGAAGTTTCACTGGCA | 92 | 61 |  |
| Acta2 | GATCAAGATCATTGCCCCTCC | GCCCGGCTTCATCGTATTCC | 122 | 61 |  |
| Tgfb1 | CCGTGGAGGGGAAATTGAGGG | AACCCGTTGATGTCCACTTGC | 90 | 61 |  |
| IL-6 | GGATTCAATGAGGAGACTTGCC | ACTCTCAAATCTGTTCTGGAGG | 93 | 61 |  |
| MMP1 | GTCACACCTCTGACATTCACC | GAGTTGTCCCGATGATCTCCC | 91 | 61 |  |
| MMP2 | CCCGGAAAAGATTGATGCGG | GTAGATCCAGTATTCATTCCCTGC | 82 | 61 |  |
| MMP9 | CTGGAGGTTCGACGTGAAGG | TAGGCTTTCTCTCGGTACTGG | 120 | 61 | ^2^ |
| SPP1 | GCCGACCAAGGAAAACTCAC | CACAGGTGATGCCTAGGAGG | 70 | 61 |  |
| PTGS2 | AATCCTTGCTGTTCCCACCC | GTCAAAAATTCCGGTGTTGAGC | 128 | 61 |  |
| TIMP1 | CACTACCTGCAGTTTTGTGGC | ATGGATAAACAGGGAAACACTGTGC | 117 | 61 |  |
| GAPDH | TGCACCACCAACTGCTTAGC | GGCATGGACTGTGGTCATGAG | 87 | 58 |  |

Supplemental Figure 1.

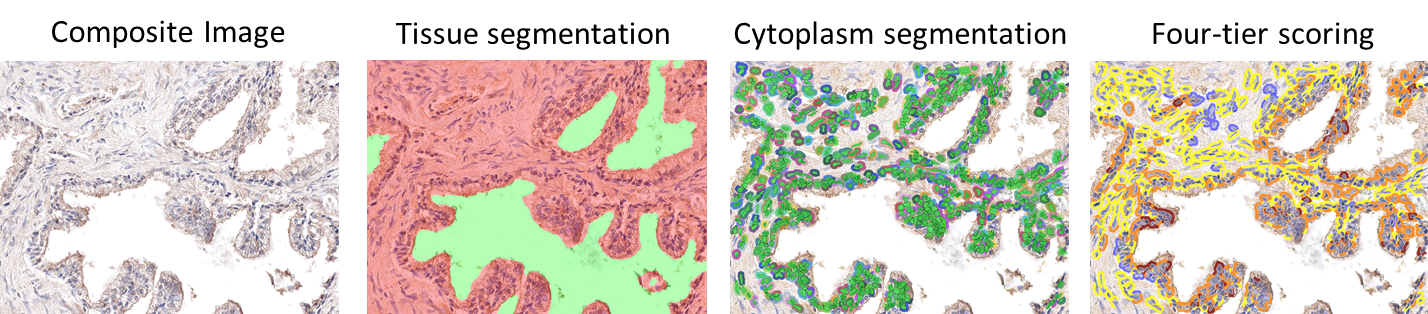

Supplemental Figure 1.: Pipeline for IHC scoring. The area of tissue was first segmented from the lumen, the nuclei were identified based on hematoxylin staining and cytoplasm was determined as the 20-pixel area surrounding the nucleus. Then tissues were scored based on 4 manually determined intensity-intervals set for batch analysis.

Supplemental Figure 2.

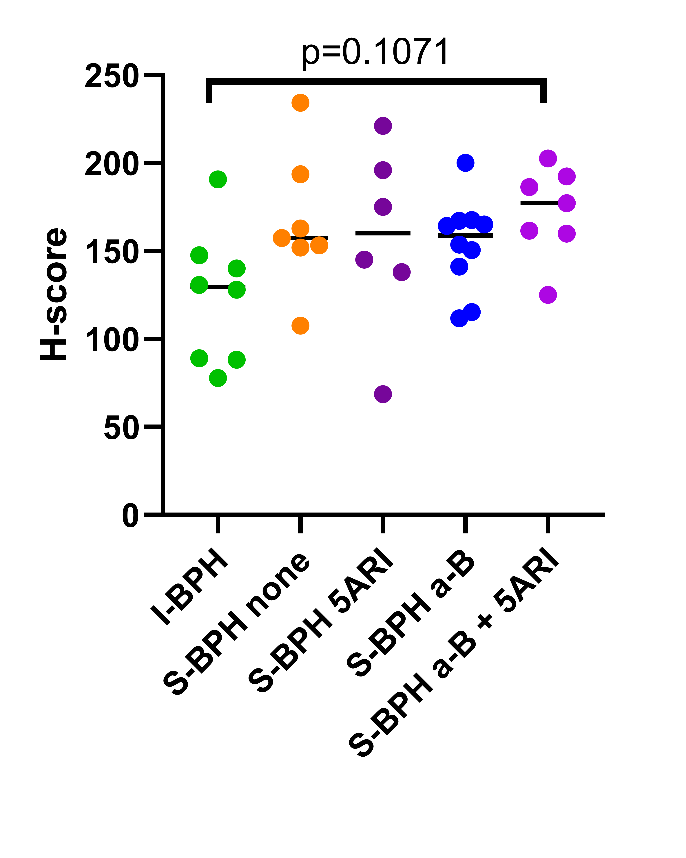

Supplemental Figure 2.: Distribution of H-scores for OPN staining across different treatment groups. 5ARI: 5α reductase inhibitors, a-B: α adrenergic antagonists.
